# Testing for pollinator recognition in multiple species of *Heliconia*

**DOI:** 10.1101/2021.04.08.438933

**Authors:** Dustin G. Gannon, Adam S. Hadley, Urs G. Kormann, F. Andrew Jones, Matthew G. Betts

## Abstract

Many plants have evolved floral traits that, in effect, filter pollinator communities and promote pollination by efficient pollinators. Most documented pollinator filter traits act to deter or reduce visitation rates by a subsect of the community of floral visitors. However, a recently described pollinator filter termed ‘pollinator recognition’ (PR) acts at a stage after a pollinator visit. PR was first documented experimentally in *Heliconia tortuosa* whereby pollen tube germination – a proxy for reproduction – was enhanced following visits from morphologically specialized pollinators, but not generalists. This is thought to promote outcrossing among plants by preferentially investing in reproduction following visits by long-ranging hummingbirds with specialized bill shapes. To date, this plant behavior has only been described in *H. tortuosa*, but, if widespread, could have important ecological implications; given declines in abundances of specialist pollinators, visits by generalists would not buffer the loss of pollination services to plants with PR. We therefore tested for PR in four taxa spread widely across the Heliconiaceae.

We corroborated previous results that visits by long-billed, but not short-billed hummingbirds increased pollen tubes in *H. tortuosa* with aviary experiments that standardized pollen quality and minimized variation in pollen quantity. Across species, we found great variation in pollen tube responses to experimental treatments. For one species (*H. rostrata*), we found increased numbers of pollen tubes in those visited by hummingbirds compared to hand pollination alone, regardless of the visiting bird’s bill morphology, indicating recognition of hummingbirds in general. In other cases, hummingbird visits decreased pollen tube counts compared to hand pollinations alone. Furthermore, our results could not substantiate any specific mechanism for pollinator recognition and highlight the need for further work on the complexities of and variability in reproductive strategies across plant taxa.

## Introduction

Pollinator filters are floral traits that, in general, manipulate animal visitation patterns presumably to promote conspecific pollen transfer or limit access to floral rewards (e.g., nectar) for poor pollinators. For example, nectar that is distasteful to some pollinators will deter them from visiting [1] and exploitation barriers, such as long corolla tubes that limit access to nectaries [2,3], may make visitation unprofitable for some animals [4,5]. These filters reduce the richness of pollinator species (or types within species) that visit a given flower species or type. However, previous work with *Heliconia tortuosa* (Heliconiaceae) documented a cryptic pollinator filter that could act at a stage after the pollinator visit to promote pollen germination and pollen tube growth based on the identity, behavior, and bill morphology of hummingbird floral visitors [6].

In single-visit aviary experiments that controlled for variation in pollen deposition and visitation rates by different pollinator species [6], the average number of pollen tubes that germinated in a style (henceforth ‘pollen tube rate’) was nearly six times greater in flowers visited by hummingbirds with bill shapes that are morphologically matched to the flowers (i.e., long, decurved bills) than in flowers visited by hummingbirds with mismatched bill shapes. Furthermore, in a separate experiment, manual nectar removal showed higher pollen tube rates than hand pollination alone. Betts et al. coined this behavior ‘pollinator recognition’ and posited that nectar removal and pollen deposition by long-billed hummingbirds provides a cue for pollen grain germination and pollen tube growth, thus reducing pollination efficiency by morphologically mismatched hummingbirds that visit and transfer pollen but cannot access the full volume of nectar at the base of the flower [6].

Betts et al. (18) speculated that pollinator recognition may be adaptive if it allows plants to invest in reproduction following visits from high-quality pollinators (those more likely to carry high-quality, outcrossed or unrelated pollen) and limit reproduction with the pollen deposited by poor pollinators (those more likely to carry low-quality pollen). Despite receiving visits from at least six hummingbird species, *H. tortuosa* specializes on long-billed hummingbirds that are highly mobile [6,7] compared to the short-billed hummingbirds which tend to defend territories and therefore move less. The mobile foraging behaviors of these birds may make them more likely to carry high-quality pollen from geographically and genetically distant sources [8], thus promoting outcrossing and genetic diversity in *H. tortuosa* populations.

We postulated that pollinator recognition may occur in other plant taxa, particularly in relatively stable tropical systems with high pollinator diversity. Determining whether this is the case is important for two reasons. First, we agree with Betts et al. [18] that filtering the short-billed, territorial hummingbirds could promote outcrossing and enhance the genetic diversity of pollen grains that reach the ovules, a hypothesis that is supported by landscape genetic studies of *H. tortuosa* populations in Costa Rica [9,10]. Given the potential for fitness benefits, pollinator recognition could be present in many related taxa and could be one means through which tight morphological matching evolves despite apparently generalized interaction networks. Indeed, hummingbird-*Heliconia* systems are often used as outstanding examples of morphological matching (e.g., [11]).

Second, if the capacity for plants to actively filter floral visitors based on morphological trait matching is widespread, this would have implications for the robustness of plant-pollinator communities under climate and anthropogenic change [12,13]. Given local extinction or reduced densities of morphologically matched pollinators, mismatched pollinators may alter their foraging behaviors to exploit newly available resources [14–17]; however, visits from mismatched pollinators would not compensate for the pollination services lost to a plant with a pollinator recognition mechanism, even if they deposit pollen at the stigma. This could increase the likelihood of coextinctions [12,18]. Thus, we sought to test for pollinator recognition in four species distributed widely across the Heliconiaceae phylogeny as a first step in assessing the generality of this cryptic pollinator filter.

## Materials and Methods

### Ethics statement

All experimental methods involving hummingbirds were approved by the Oregon State University Animal Care and Use Committee (Animal Care and Use Permit 5020) and all international research guidelines and practices were followed.

### Study species

Heliconiaceae is a monogeneric family consisting of an estimated 200-250 species which radiated rapidly c.a. 39-24 million years ago [19]. *Heliconia* species are rhizomatous perennial herbs distributed widely throughout the Neotropics and on some South Pacific islands. Flowers are situated within showy bracts and composed of six tepals, five of which are fused to create a cylindrical perianth, the sixth peels back upon anthesis. A defining feature of the Heliconiaceae is a staminode (modified stamen) that partially covers the opening to the nectar chamber at the base of the perianth, which may need to be moved by a visiting animal when they extract the nectar reward (though the mechanics of this have not be studied in detail). Flowers of the Heliconiaceae last a single day from anthesis to dehiscence.

We targeted species that were common in the living collection at the Organization of Tropical Studies Las Cruces Biological Station, Puntarenas Province, Coto Brus, Costa Rica, (8° 47’ 7” N, 82° 57’ 32” W) and could be found naturally or in ornamental gardens in the area. We required that plants were setting seed when left unmanipulated, indicating that a viable pollen source existed in the area, since previous work on mating systems in *Heliconia* suggests that the hermaphroditic flowers of many species are self-incompatible to partially self-compatible, but largely not selfing [6,9,20–23]. Furthermore, we required that wild, native hummingbirds could be seen visiting the flowers of each target species in camera trap data [24] or during observation, indicating that wild-caught hummingbirds would visit and drink from the flowers inside aviaries despite the fact that many plant species in the collection are not native to Costa Rica. The plant species that met these criteria included *H. hirsuta*, which is native to South America and Trinidad [25], *H. rostrata*, native to western South America [25] but a common ornamental throughout the tropics, and *H. wagneriana*, native to Costa Rica and Panama [26]. Furthermore, because so little is known of this unusual plant behavior, we also sought to replicate the results of the original study in the native *H. tortuosa*, an exercise rarely undertaken in experimental ecology [27].

We selected two hummingbird species with different bill morphologies and foraging behaviors as “treatments” in order to accentuate differences in morphological matching to and nectar depletion from the range of flower shapes exemplified by the four *Heliconia* species (Fig 1). Green Hermit Hummingbirds (*Phaethornis guy*; GREH) are common traplining hummingbirds in the region with long (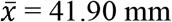, *s_x_*= 1.52 mm), moderately decurved bills (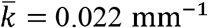, *s_k_* = 0.004 mm^-1^, *n* = 27 birds of mixed sex, where *k_i_*, is the curvature of the *i*^th^ bill measured as the inverse of the radius of the arc of the bill – see Temeles et al. [3]). Rufous-tailed Hummingbirds (*Amazilia tzacatl*; RTAH) are common territorial hummingbirds with short (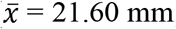, *s_x_* = 1.55 mm, *n* = 14 birds of mixed sex), slightly decurved bills (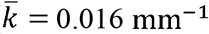, *s_k_* = 0.002 mm^-1^; Fig 1).

**Fig 1.**
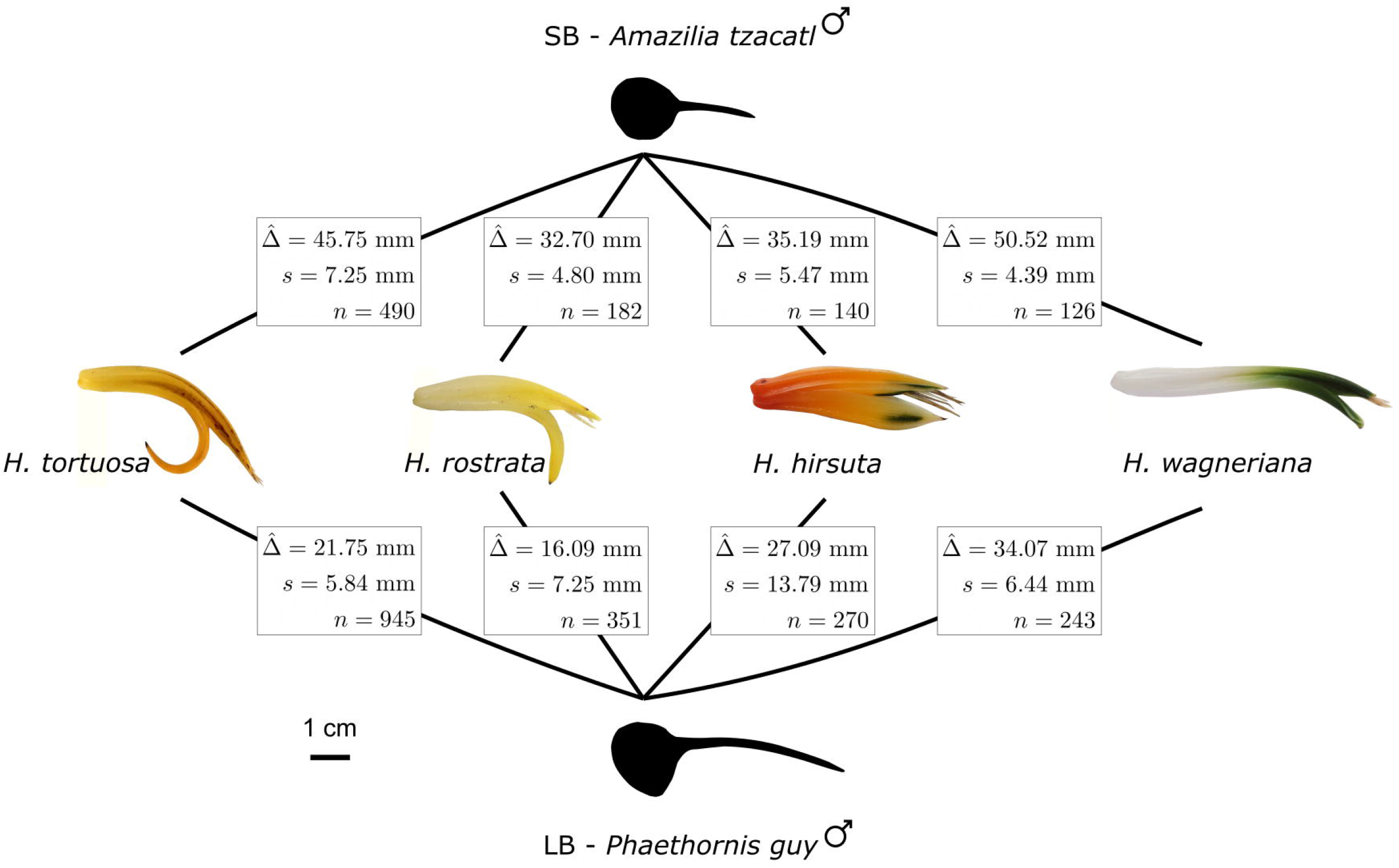
Morphological mismatch between the hummingbird and *Heliconia* species used in experiments. The average mismatch 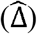 between a plant and hummingbird species was measured as the Euclidean distance between a flower and a bird’s bill in the 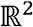 trait space, where one axis was the total length of a bill or flower (mm) and the other was the radius of the arc along the outside edge of the flower or bill (mm). We then computed the mean 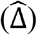 and standard deviation (*s*) of the distances between each bill-flower pair, where *n* is the number of pairwise comparisons.

Assuming that morphologically matched floral visitors increase the numbers of successful pollen tubes for all *Heliconia* species, we predicted the following: 1) For *H. wagneriana* and *H. tortuosa*, we would predict greater numbers of pollen tubes in flowers visited by green hermit hummingbirds compared to rufous-tailed hummingbirds due to long and curved flowers (Fig 1). 2) Because both *H. hirsuta* and *H. rostrata* have shorter, straighter flowers and both hummingbird bill shapes approximate the shape of the flowers well (Fig 1), we would not predict a large difference in the number of pollen tubes between flowers visited by green hermits and those visited by rufous-tailed hummingbirds. We therefore used hand pollinations as a control treatment in all experiments since hand pollinations do not replicate the physical characteristics of a visit by a morphologically matched pollinator aside from pollen deposition. Hence, we would predict the fewest pollen tubes in flowers pollinated by hand for all plant species. Furthermore, this helped us control for potentially low genetic diversity in the pollen pool since the control flowers (hand pollination only) and the treatment flowers (hand pollination followed by a visit from a pollen-free hummingbird) both received pollen by hand from the same donor.

### Aviary experiments

To test whether pollen germination and tube growth is dependent on interactions with morphologically matched floral visitors, we conducted 110 single-visit experiments (*n* = 214 flowers from 54 plants; see Table 1 for the number of replicates per treatment) with captive hummingbirds inside portable aviaries. The aviaries measured 2 meters tall and one meter on a side and could be quickly assembled around live plants (S1 File). In these experiments, we used only virgin flowers that had been covered with mesh bags prior to anthesis in order to preclude pollination by free-ranging pollinators. Flowers were not emasculated, however, due to low numbers of pollen tubes in emasculated flowers in earlier experiments with plants in aviaries and in natural settings (M. G. Betts and A. S. Hadley, *unpublished data*).

**Table 1:** Sample sizes for individual plants (grouping factor) and flowers (experimental units) for each species × treatment combination.

|  | Hand<br>pollination<br>(HP)* | Hand<br>pollination<br>+ short-billed bird<br>(SB) | Hand<br>pollination +<br>long-billed<br>bird (LB) | Hand<br>pollination<br>(HP)** | Hand<br>pollination<br>+ pipette<br>(BM) | Hand<br>pollination<br>+ nectar<br>extraction<br>(HPNE) | Nectar<br>extraction<br>+ hand<br>pollination<br>(NEHP) |
| --- | --- | --- | --- | --- | --- | --- | --- |
| ***H. hirsuta*** |  |  |  |  |  |  |  |
| Plants | 7 | 4 | 5 | 0 | 0 | 0 | 0 |
| Flowers | 11 | 5 | 7 | 0 | 0 | 0 | 0 |
| **<u>H. rostrata</u>** |  |  |  |  |  |  |  |
| Plants | 19 | 13 | 10 | 13 | 7 | 15 | 5 |
| Flowers | 39 | 25 | 19 | 24 | 8 | 38 | 11 |
| **<u>H. tortuosa</u>** |  |  |  |  |  |  |  |
| Plants | 10 | 8 | 6 | 10 | 7 | 10 | 4 |
| Flowers | 16 | 10 | 11 | 27 | 7 | 36 | 5 |
| **<u>H. wagneriana</u>** |  |  |  |  |  |  |  |
| Plants | 12 | 6 | 11 | 11 | 7 | 8 | 4 |
| Flowers | 31 | 17 | 23 | 17 | 7 | 12 | 10 |

\*Hand pollination controls for aviary experiments
\*\*Hand pollination controls for nectar removal experiments

We selected inflorescences based on the availability of two virgin flowers and erected the aviary around the whole plant. We then randomly assigned one of the flowers as a control flower that received hand-pollination but no visit from a bird (HP treatment). The remaining flower was hand-pollinated with pollen from the same donor flower, then allowed a visit by either a pollen-free short-billed hummingbird (SB treatment; *n* = 14 *A. tzacatl* individuals used in experiments) or a pollen free long-billed hummingbird (LB treatment; *n* = 12 *P. guy* individuals used in experiments). To ensure the birds were free of pollen before the visit to the focal flower, we cleaned them using a soft paint brush and damp cotton swab under 20x magnification prior to releasing them into the aviary. Thus, flowers were the experimental units and individual plants were treated as a blocking effect to account for potential dependence among measurements on flowers from the same plant. Where possible, plants received all treatments (often on separate days).

By hand-pollinating all flowers using pollen from an arbitrarily selected pollen donor, we were able to control for differences in the quality of pollen delivered by the different pollinator species. Indeed, we could not perfectly standardize the quantity of pollen grains at the stigmatic surface because the size of *Heliconia* pollen grains makes it impractical to quantify the number of grains in the field; however, we attempted to reduce variation in the quantity of pollen available to the flowers by having the same experimenter apply pollen in an even layer across the stigmatic surface with a toothpick under 20x magnification for every flower.

After the hummingbird visited the treatment flower (evidenced by bill insertion and a clear attempt to feed from the flower), we terminated the experiment and checked the stigma again to ensure that pollen was still present and in an even layer on the stigmatic surface before again covering the flowers with mesh bags. All flowers were collected the following day, the styles removed and preserved in formalin acetyl-acid, and scored for pollen tubes using epiflorescence microscopy [6,20] (see S1 File for more information). All aviary experiments were conducted during the 2018 and 2019 dry seasons (Feb-Mar).

### Tests for a mechanism

We conducted additional experiments to test hypotheses of the mechanism of pollinator recognition. Betts et al. [6] found increased pollen tube rates in flowers from which nectar was removed compared to hand pollination alone. As an independent test of whether nectar removal provides a cue to which plants respond, we manually extracted nectar from flowers of three of the four species (*H. hirsuta* did not produce flowers regularly enough to conduct the full suite of experiments) and compared pollen tube rates to control flowers that were hand-pollinated on the same day. Alternatively, it is possible that the long-billed hummingbirds trigger a mechanical cue [28] when they insert their bills into the flower. To test whether we could induce an increase in pollen tube success rates using a mechanical stimulus, we molded a pipette tip to match the curvature of the focal flower. We then inserted the pipette tip as a hummingbird would insert its bill but did not remove any nectar. Finally, because we were unable to perfectly replicate the timing of events in a natural pollinator visit in which nectar removal and pollen deposition happen concurrently, we conducted some experiments in which we hand pollinated before manually removing nectar and some in which we hand pollinated after removing nectar. Differences in these pollen tube rates may indicate the importance of the timing of pollen transfer and nectar removal or bill insertion (see S1 File for more detail).

### Statistical methods

We separately analyzed pollen tube count data from each set of experiments (i.e., aviary experiments as one dataset and nectar removal experiments as a second dataset). We fit generalized linear mixed models (GLMMs) to both sets of data using the R package lme4 [29,30] (see S1 File for full model specification). Our experiments were factorial with each plant species receiving each treatment (4 plant species × 3 treatments = 12 means to estimate for the aviary data; 3 plant species × 4 treatments = 12 means to estimate for the nectar removal data). Furthermore, we included random effects for each plant (nested within plant species) to account for potentially correlated observations that could arise from scoring pollen tubes in multiple flowers from the same plant (since individual plants received more than one treatment).

Below, we report mean pollen tube counts per style estimated from the model fit for a given treatment and plant species as 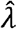 with a subscript indicating the treatment and plant species. We use *HP, SB, LB, HPNE, NEHP*, and *BM* to indicate the treatment (Table 1). Treatment codes are as follows: *HP* identifies the hand-pollinated control flowers; *SB* indicates the treatment in which we hand pollinated flowers, then allowed a clean, rufous-tailed hummingbird (short, straight bill) to visit; *LB* indicates the treatment in which we hand pollinated flowers, then allowed a pollen-free green hermit hummingbird (long bill) to visit; *HPNE* identifies the treatment in which we hand pollinated the flowers then manually removed the nectar; *NEHP* identifies the treatment in which we hand pollinated the flowers after removing nectar; and *BM* identifies the treatment in which we inserted a pipette tip but did not attempt to remove nectar. We use the letters *h, r, t*, and *w* to identify *H. hirstuta, H. rostrata, H. tortuosa*, and *H. wagneriana* in the subscripts (respectively). We additionally report differences between treatments as the fold change in pollen tube rates and use the notation 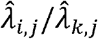 for the fold change between treatments *i* and *k* for plant species *j*. This is a natural way in which to report differences between treatments given the log link function. Finally, we estimated ninety-five percent confidence intervals from 1,000 bootstrapped mean predictions [29]. Confidence intervals for the true parameters follow each estimate and are presented in square brackets.

## Results

When we compared pollen tube counts in hand-pollinated control flowers (HP) and those that were visited by a pollen-free, morphologically matched hummingbird, we found evidence that a visit by a matched hummingbird increased pollen tube rates for *H. tortuosa* over hand pollination alone (Fig 2). Pollen tube rates were 6.91 [3.56, 43.49] (estimate and 95% confidence interval) times greater following visits from long-billed hummingbirds compared to the control treatments with only hand pollination (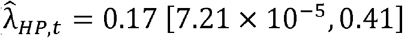 tubes per style; 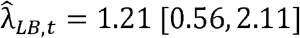 tubes per style). However, short-billed hummingbird visits did not boost pollen tube rates for *H. tortuosa* much above the hand-pollinated controls (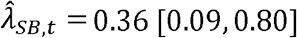 tubes per style; Fig 2). Thus, the pollen tube rate in flowers visited by morphologically matched hummingbirds was greater (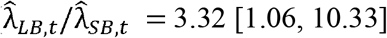 times greater) than the pollen tube rate in flowers visited by morphologically mismatched hummingbirds (Figs 1&2).

**Fig 2.**
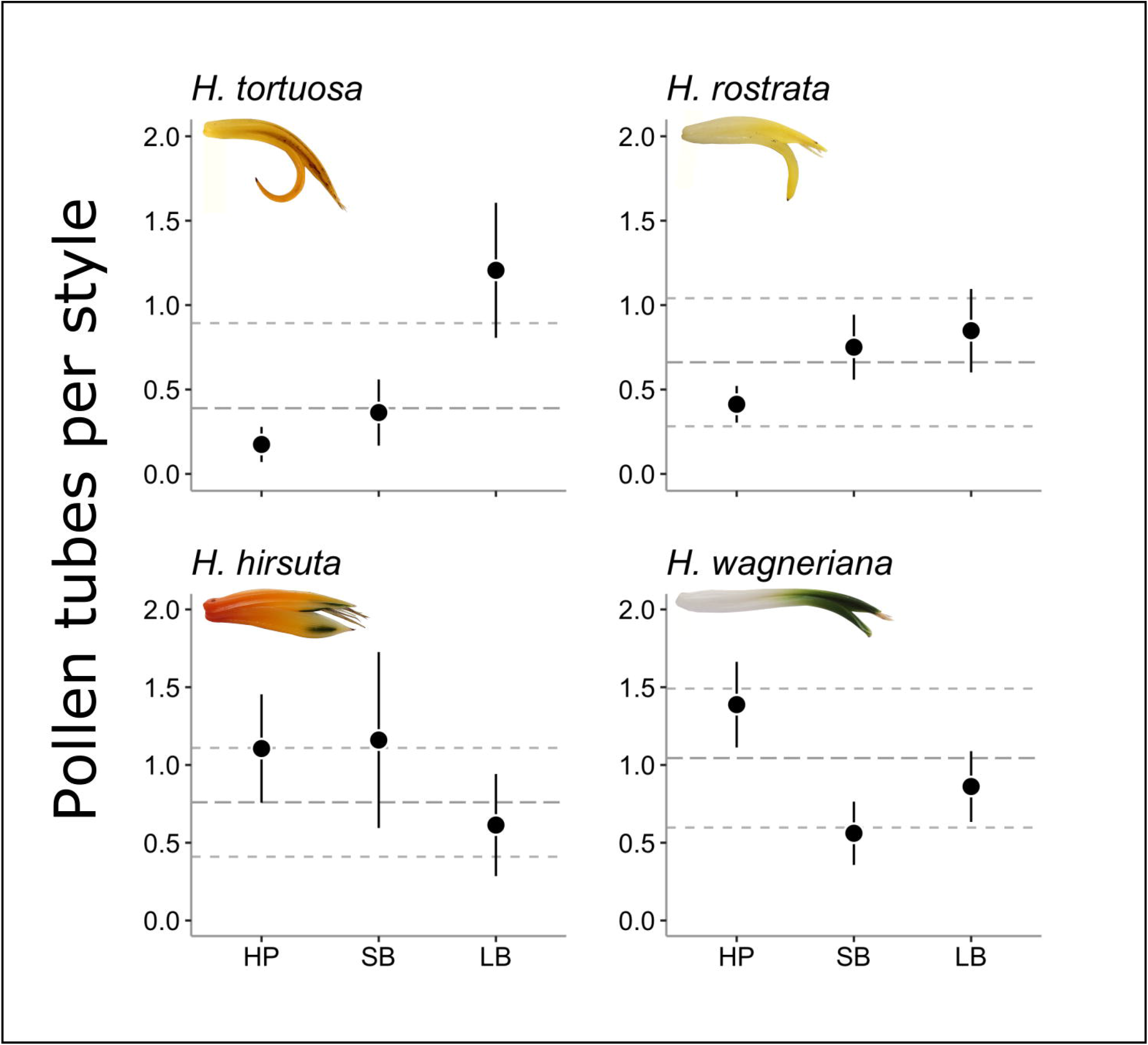
Pollen tube rates in flowers that received hand pollination only (HP) and those visited by a pollen-free hummingbird following hand pollination. Flowers were visited by either short-billed (SB) rufous-tailed hummingbirds (*Amazilia tzacatl*) or long-billed (LB) green hermit hummingbirds (*Phaethornis guy*). Error bars show the estimate plus or minus bootstrapped standard errors. The grey, horizontal dashed lines show estimates (±*SE*) of the pollen tube rates in flowers left open to free ranging pollinators.

In *H. rostrata* flowers, pollen tube rates in those visited by hummingbirds were greater than hand pollination alone regardless of the bird species used in experiments. The estimated rates were similar in flowers visited by long-billed hummingbirds and those visited by short-billed hummingbirds, but were approximately double the rate in hand pollinated controls (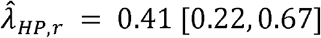 tubes per style; 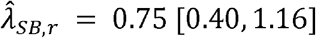 tubes per style; 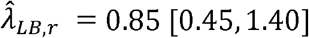 tubes per style; Fig 2).

For *H. hirsuta* and *H. wagneriana*, single visits from cleaned hummingbirds did not enhance pollen tube success rates above hand pollination alone (Fig 2), and pollen tube rates in *H. wagneriana* flowers that were visited by clean birds were actually reduced. The number of pollen tubes per style in *H. wagneriana* flowers visited by green hermits were a little more than half those of hand pollinations and short-billed hummingbird visits yielded pollen tube rates less than half of hand pollination treatments (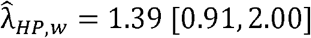 tubes per style; 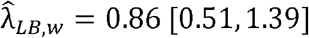 tubes per style; 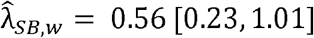 tubes per style; Fig 2).

When we experimentally removed nectar using pipette tips to test the hypothesis that differential nectar removal may be the mechanism for pollinator recognition, the effect of nectar removal on pollen tube rates for *H. rostrata* and *H. tortuosa* was negligible (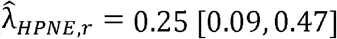 tubes per style; 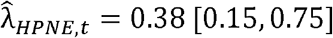 tubes per style; Fig 3). We did find weak evidence that our nectar removal treatments had a positive effect on pollen tube germination relative to hand-pollinations alone for *H. wagneriana* (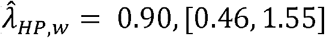 tubes per style in the nectar removal data; 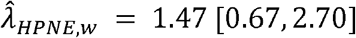 tubes per style; Fig 3), but this effect is driven in part by a small number of influential observations (>5 pollen tubes found in 2 styles), and disappears after removing them, so we caution readers in their interpretation of this result. Furthermore, we achieved higher pollen tube rates with hand pollination alone during the aviary experiments (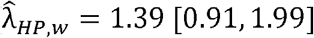 tubes per style from aviary experiments) than in the nectar removal experiments.

**Fig 3.**
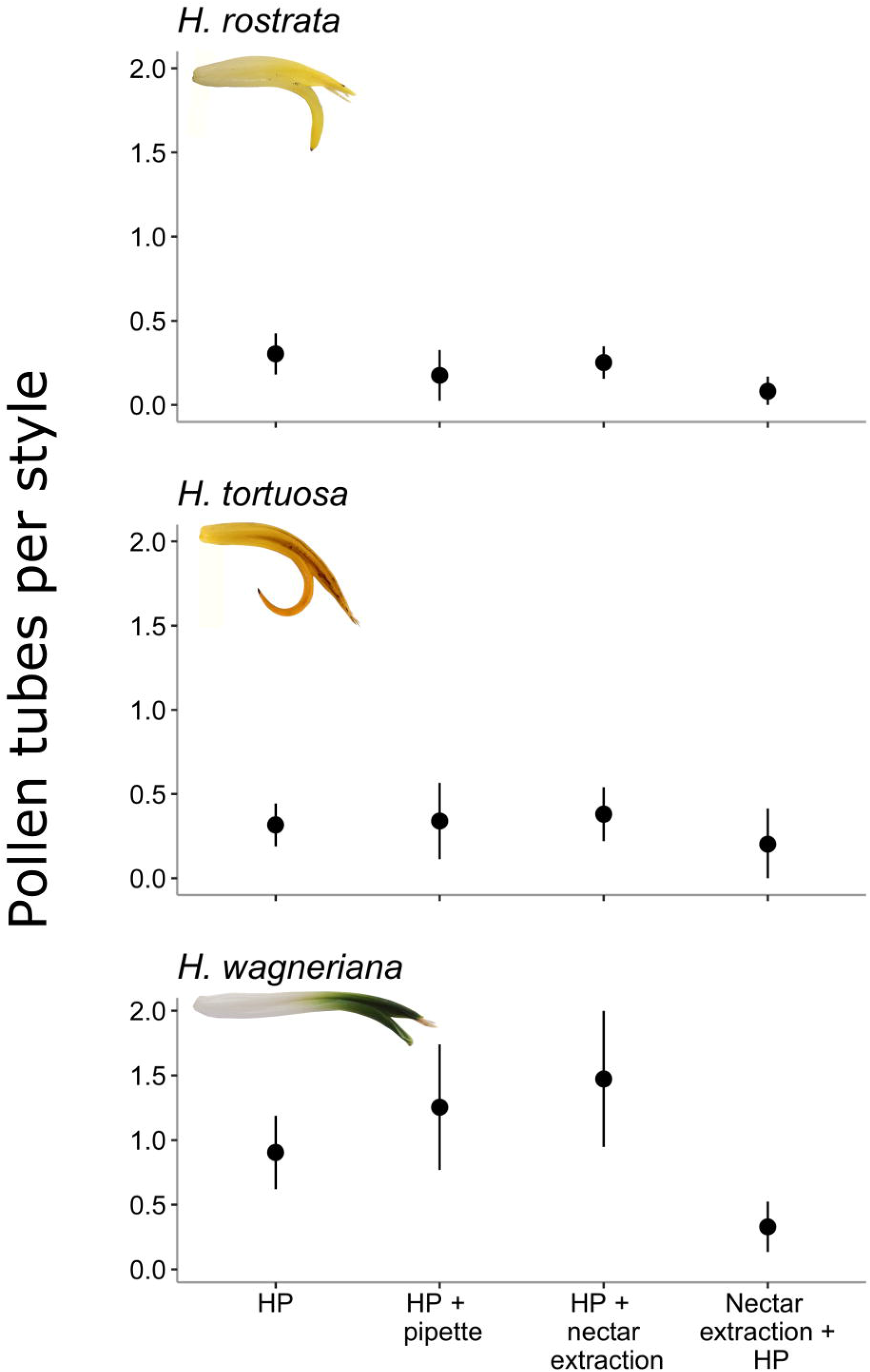
Results from experimental tests of the effect of nectar depletion on pollen tube rates. We used hand-pollination (HP) as a control treatment and compared pollen tube rates in flowers that received the control treatment to those in flowers that received out-cross pollen by hand either before (HP + nectar extraction) or after (Nectar extraction + HP) manual removal of the nectar in the flower. As a test of whether pollen germination success could be boosted after the mechanical stimulus of a hummingbird inserting its bill to drink from the flower, we tested for an effect of pipette insertion without removing any nectar (HP + pipette). Error bars show estimates of the mean plus or minus bootstrapped standard errors.

Inserting a hummingbird bill mimic (i.e., pipette tip) into flowers as a mechanical signal without removing nectar also did not induce substantially higher pollen tube rates in any of the tested species (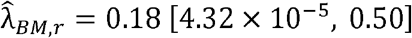 tubes per style; 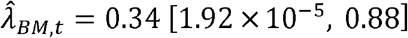 tubes per style; 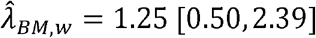 tubes per style; Fig 3), and, in all cases, hand pollinating flowers after removing nectar resulted in the fewest pollen tubes per style out of all treatments, generally about half of hand pollination alone (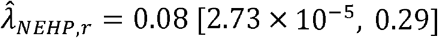 tubes per style; 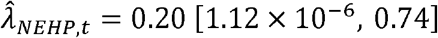 tubes per style; 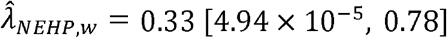 tubes per style; Fig 3).

## Discussion

We tested whether pollinator recognition, a plant behavior first described in *Heliconia toruosa* [6] in which plants may discriminate among floral visitors based on morphological matching to the flower, may be present in more than one species of *Heliconia*. Our results substantiate the findings of Betts et al. [6] that the number of successful pollen tubes in *H. tortuosa* flowers is greater in those visited by morphologically matched hummingbirds compared to those visited by mismatched birds after experimentally standardizing pollen quality, visit frequency, and minimizing variation in pollen quantity. However, we did not find strong support for any of the hypothesized mechanisms of recognition and found different responses to our aviary treatments in each of the four *Heliconia* species we tested, highlighting the poorly understood complexities of pollination in *Heliconia* plants.

The results from our aviary experiments with *H. tortuosa* are highly congruent with those of Betts et al. [6] and support that plants specialize on long-billed, traplining hummingbirds.

Pollinator recognition in *H. tortuosa* is likely an additional pollinator filter acting in conjunction with morphological barriers that often result in only imperfect resource partitioning by floral visitors [4,31–33]. However, this could have strong ecological implications if disturbances reduce the abundance of morphologically matched hummingbirds and the rewiring of the remaining community of floral visitors fails to buffer the loss of pollination services to *H. tortuosa* plants. In forest fragments around Coto Brus, mismatched hummingbirds account for c.a. 10% of honest visits (those in which the visitor contacts the reproductive organs of the flower) to *H. tortuosa* (K. Leimburger, *unpublished data*), but this proportion likely increases in isolated fragments where morphologically matched hummingbirds are less common [34–36]. The combination of fragmentation and pollinator recognition therefore has a strong ecological impact on *H. tortuosa* populations, which show reduced seed sets in fragmented forests compared to those in continuous forest [37], presumably due to a paucity of morphologically matched hummingbirds.

In *H. rostrata* styles, visits from clean hummingbirds to hand-pollinated flowers also increased pollen tube rates, but regardless of the bird species. The estimated pollen tube rate in *H. rostrata* flowers that were visited by long-billed hummingbirds was nearly identical to that estimated for flowers visited by short-billed hummingbirds (Fig 1). However, given the relatively short, straight corolla of *H. rostrata* (Fig 1), both hummingbird species we used for experiments were able to achieve strong morphological matches and might not be expected to differ in their visitation characteristics, such as nectar consumption. We did not destructively sample flowers after hummingbird visits to measure the nectar remaining, but both species of birds can be seen drinking from *H. rostrata* flowers in recorded videos (S1 and S2 videos).

In Peru, seven hummingbird species of various sizes and with various bill shapes have been observed visiting *H. rostrata*, but nothing is known of their pollination efficiencies [38]. Based on our results showing increased pollen tube rates in bird-visited flowers compared to hand pollination, we posit that *H. rostrata* could filter visits from animals without complementary morphologies, such as insects. This idea is supported by data from Janeček et al. [23] who recorded olive sunbirds (*Cyanomitra olivacea*) and Camaroon sunbirds (*Cyanomitra oritis*) visiting *H. rostrata* flowers in South Africa where it has been introduced. These authors found that *H. rostrata* flowers left open to visits from sunbirds had extremely low pollen tube rates, as did hand-pollinated flowers. Further work in the native range of *H. rostrata* is necessary to determine if *H. rostrata* cryptically filters the community of floral visitors.

We did not detect strong differences among treatments in *H. hirsuta* flowers, which all showed pollen tube rates in the range of open pollination pollen tube rates (grey lines in Fig 2). Interestingly, *H. hirsuta* flowers visited by long-billed hummingbirds following hand pollination showed the lowest pollen tube rate. Similarly, we found reduced pollen tube rates in *H. wagneriana* flowers that were visited by hummingbirds relative to hand-pollinations alone (Fig 2). While the mechanisms underlying these results in *H. wagneriana* and *H. hirsuta* flowers remain unclear, we identified one way in which these species differ from the others that could produce this result. Gannon et al. [24] discovered that *H. wagneriana* plants have a mechanism for keeping the anthers protected within the perianth and then rapidly extending them as a hummingbird visits. This is thought to protect pollen from desiccation and/or increase pollen transfer to pollinators during the first visit. Once exposed, however, pollen grains desiccate relatively quickly, and often fail to adhere to the stigmatic surface. While *H. hirsuta* anthers protrude from the flower upon anthesis, pollen also appeared to dry quickly and often failed to cling to the stigmatic surface. This may make the pollen grains of *H. wagneriana* and *H. hirsuta* easy to dislodge. While we checked that pollen was still present on the stigma after a bird visited, the size of *Heliconia* pollen makes exact quantification in the field infeasible. Thus, it is possible that reduced outcross pollen loads after the birds visited resulted in reduced pollen tube counts relative to hand pollination alone.

Using camera traps, Gannon et al. [24] found that c.a. 97% of the visits to open *H. wagneriana* flowers around Las Cruces were by traplining species with morphologically matched bill shapes. Similarly, Snow and Snow [39] report only green hermit and rufous-breasted hermit (*Glaucis hirsutus*) visitors at *H. hirsuta* flowers in Trinidad (part of its native range), both of which have well-matched bills. Given a relatively specialized set of floral visitors to these two species, it seems unlikely that any cryptic recognition mechanism would adaptively filter the community of floral visitors.

### The mechanism of pollinator recognition

Previous work demonstrated increased pollen tube rates with manual nectar extraction treatments compared to hand pollinations alone in *H. tortuosa* [6]. Betts et al. [18] hypothesized that, because birds with well-matched bill morphologies can drain the nectar chamber but those with mismatched bills often cannot [5,6], nectar removal could provide a cue to which plants respond to promote successful pollen tube growth. We were unable to reproduce an increase in pollen tube rates using manual nectar removal techniques, despite the use of identical nectar removal methods (aside from the experimenter). While we found increased pollen tube rates in *H. wagneriana* flowers following manual nectar removal (Fig 3), this result is tenuous because it is in part driven by a few influential observations. Furthermore, the pollen tube counts in the flowers from which we removed nectar are approximately equal to the pollen tube counts in our hand pollinated flowers from the aviary experiments (Figs 2 and 3).

In summary, the evidence that nectar removal provides the cue for pollinator recognition in *H. tortuosa* is equivocal and further experiments are necessary to verify nectar removal or establish a new mechanism. We refrain from speculating on alternative mechanisms of recognition, but instead suggest follow-up experiments that could help to disentangle the apparent complexities of this system: 1) continued aviary experiments with Heliconia species in in their native ranges with native floral visitors; 2) comparative studies of the mechanics of visits to *H. tortuosa* flowers by different hummingbird species using high-speed videography to identify potential mechanisms for higher pollen tube rates by morphologically matched species; 3) enhanced experimental techniques that can increase the consistency of manual nectar extraction and test alternative potential mechanisms linked to morphological matching (including, but not limited to, nectar removal, pollen deposition/removal, and other aspects of visit mechanics).

## Conclusions

Our results help to highlight the complexities of pollination and pollen germination success. Despite our detailed and manipulative experiments, we found great variability in the number of germinated pollen tubes depending on the floral visitor and plant species. We therefore add to calls for more detailed and manipulative experiments to assess realized pollination network structure and vulnerability to disturbance that take into account single-visit pollination efficiency [40,41]. Furthermore, our results highlight the necessity and opportunity for further study of pollinator recognition in *Heliconia*. Further research into the mechanism of recognition in *H. tortuosa* and the generality of pollinator recognition in the Heliconiaceae is warranted. Notably, Pedersen and Kress [21] report a c.a. four-fold increase in pollen tube rates in *Heliconia paka* flowers that were visited by honeyeaters compared to those pollinated by hand. These results would be consistent with what we would predict for *H. paka* given a pollinator recognition mechanism. More generally, Young and Young [42] report that hand-pollinated flowers had reduced reproductive output compared to open-pollinated flowers for 17 of 52 plant species from highly divergent lineages. We know of no follow-up experiments with these or related taxa, but we urge others to conduct similar experiments to those presented here to examine the potential for cryptic specialization in other pollination systems.

## Acknowledgements

We thank C. Dowd, E. Sandi, M. Atencio, C. Tortorelli, and G. Doyle for invaluable assistance with field experiments, and the staff of the Las Cruces Biological Station for maintaining an excellent location in which to conduct research. We thank N. Waser, R. Sargent, J. Lau, and seven anonymous reviewers for comments on previous versions of the manuscript.

## Data accessibility

Data from pollination experiments and all R code necessary to reproduce the results can be found on a public Github repository (https://github.com/Dusty-Gannon/PR-in-Heliconia).

## Author contributions

M.G.B., A.S.H., and D.G.G. designed experiments. D.G.G. analyzed the data and wrote the original version of the manuscript. All authors contributed to data collection and critical review of the manuscript.

## Supporting information

**S1 video. Green hermit visit to *H. rostrata*.** Green hermit hummingbird (*Phaethornis guy*) visiting and drinking nectar from a *Heliconia rostrata* flower.

**S2 video. Rufous-tailed hummingbird visit to *H. rostrata*.** Rufous-tailed hummingbird (*Amazilia tzacatl*) visiting and drinking nectar from a *Heliconia rostrata* flower.

**S1 File. Supplementary methods.** This file contains additional information on the field, laboratory, and statistical methods.

