## Supplemental methods for "Testing for pollinator recognition in multiple species of *Heliconia*"

### Testing for pollinator recognition in multiple species of *Heliconia*: Supplementary material

#### Contents

**Experimental methods** 1

Aviary experiments . . . 1

Nectar removal experiments . . . 2

**Laboratory methods** 4

**Trait (mis)matching** 4

**Statistical methods** 4

**References** 6

### Experimental methods

#### Aviary experiments

In identifying focal *Heliconia* species, we first identified species which were likely to have multiple individual plants (rather than multiple clones) in the Las Cruces Biological Station living collection and the surrounding area based on collection records and surveys of the area. Because *Heliconia* are self-incompatible to partially self-compatible (W. J. Kress 1983; Pedersen and Kress 1999; Janeček et al. 2020; Betts, Hadley, and Kress 2015), we sought to maximize the genetic diversity of the pollen pool in order to limit the possibility of failing to detect a difference among pollination treatments simply due to a lack of compatible pollen. As an additional confirmation that compatible pollen was available in the area, we required that plants could be seen setting fruit. Indeed, these measures did not eliminate the possibility of non-detection due to poor quality pollen since we do not know the parentage of the plants used in the experiments, but this should not inflate the chances of erroneously detecting pollinator recognition since pollen quality was held constant across control flowers and treatment flowers.

Prior to anthesis, all flowers were covered with a mesh bag in order to preclude visits from free ranging pollinators. Each flower was hand pollinated by the same experimenter (author D.G. Gannon), using pollen sourced from plants located at least five meters away from the focal plant in order to reduce the chances of applying self or related pollen to the stigma (mean distance to donor ± 1 sd: $\bar{x}$ = 233m ± 223m). We gently separated styles from the stamens using forceps and cleaned all pollen from the stigma with a cotton swab under 20x magnification. We then adhered pollen to the stigmatic surface by scraping the pollen from an anther of the donor flower with a toothpick and touching the toothpick to the stigma of the focal flower. We checked that pollen adhered to the stigmatic surface and that pollen was dispersed across the stigma in a relatively even layer using a 20x hand lens. As mentioned in the main text, quantification of pollen grains on the stigma in the field is not feasible due to the size of the pollen grains, but we attempted to minimize variation in the quantity of pollen applied across treatments and replicates.

Aviary experiments began by locating a bagged inflorescence with two mature flowers. We hand-pollinated each flower using the methods described above, then assigned flowers to a treatment at random, one flower assigned as a control (hand-pollination only) and the other to be visited by a hummingbird (long-billed or short-billed). The control flower was covered with a red paper sleeve to block access. We erected small, portable aviaries (1m x 1m x 2m) to enclose focal plants (Fig S1). Aviaries were constructed from sewn shade cloth, a one-inch PVC hoop at the top, and bamboo legs that could be embedded in the ground.

We used two common hummingbird species in our aviary experiments which represent guilds of short-billed and long-billed hummingbirds: rufous-tailed hummingbirds (*Amazilia tzacatl*), and green hermit hummingbirds (*Phaethornis guy*), respectively (Fig 1, main text). We captured hummingbirds using standard mist-netting procedures (OSU ACUP 5020), running nets from 0600 to 1000 hours. Males of focal bird species were placed in cloth bags for transport to the aviary, and all non-target species were immediately released. On some occasions, male hummingbirds birds were housed in a 2 m × 2 m × 1.5 m aviary for up to seven days to facilitate data collection (OSU ACUP 5020). This became necessary as capture rates decreased in the area of the experiments.

To ensure that hummingbirds did not contribute additional pollen to the stigmatic surface of the focal flower, we cleaned hummingbirds of all pollen under 20x magnification using damp cotton swabs and a photographer's brush before releasing them into the aviary. Hummingbirds were provided a perch inside the aviary and supplemental sugar water (20% sucrose by mass) if the hummingbird did not visit the focal flower after 30 minutes post-entry. At 60 minutes, if the hummingbird had not visited the focal flower, we terminated the experiment and the bird was either released or fed sugar water and moved to another experiment. We conducted all pollination experiments between the hours of 0600 and 1100, as *Heliconia* pollen viability and/or stigmatic receptivity may be in question later in the day (Dafni and Firmage 2000; Hedhly, Hormaza, and Herrero 2003; Schleuning et al. 2011). Flowers were labeled, covered with mesh bags to ensure no additional pollinator visits, and collected the following day after abscission.

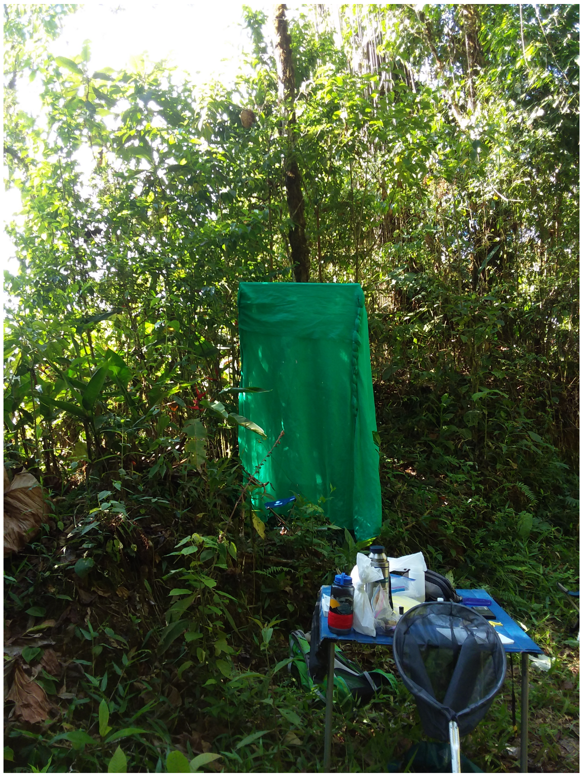

Figure S1: Portable aviary assembled around a live inflorescence.

#### Nectar removal experiments

To test whether nectar removal may provide a mechanism conferring pollinator recognition in *Heliconia*, we manually extracted nectar from flowers of three of the six focal species based on flower availability. We again included at least one control flower for each day and for each species on which conducted the experiments. We extracted nectar from flowers using a 20 $\mu$L micropipette tip bent to match the curvature of the flower. To the back of the pipette tip, we fit a length of 0.5 mm plastic tubing, connected to a 20 mL syringe with which we created suction. We measured the volume of liquid removed with a glass capillary tube, then dispensed it onto a temperature-calibrated hand refractometer to verify that the liquid was likely nectar produced by the flower. The minimum Brix index observed in an independent dataset of Heliconia nectar was c.a. 14.1% (K. Leimberger *unpublished data*). Thus, we recorded instances in which the sugar concentration measured below this level.

An alternative stimulus to which plants could respond is the mechanical action of a hummingbird inserting its bill into the perianth to access the nectar. To test whether we could induce pollen tube growth by the mechanical stimulus of inserting a morphologically matched object into the flower, we conducted experiments in which we simply inserted the pipette tip, then removed it without extracting any nectar. Nectar volumes moving up into the pipette tip due to capillary action were, except for in a few cases, not measurable. This experiment is therefore not confounded with the nectar removal experiments.

Indeed, our nectar removal treatments could not completely mimic the way in which birds interact with a flower, since pollinators deposit pollen at the same time nectar is removed. In our experiments, we could only complete these tasks in sequence. We therefore conducted some nectar removal experiments in which we hand pollinated the flower after removing nectar. These experiments allowed us to assess whether timing of nectar removal relative to pollen deposition could be important to pollen germination success.

### Laboratory methods

We collected styles the following day (after abscission) and fixed them in formalin acetyl acid for at least 72 hours before transferring them to 70% ethanol for transport back to Oregon State University. We stained pollen tubes using aniline blue dye in a buffer of $KHPO_4$ preceded by four wash steps: 24 hours in $dH_2O$, 24 hours in 5M NaOH to soften the tissues, followed by two 24-hour $dH_2O$ washes (Kress 1983; Betts, Hadley, and Kress 2015). We mounted styles on microscope slides and scored pollen tubes by counting the maximum number of pollen tubes found in any given cross-section of the style using an epi-fluorescence microscope.

### Trait (mis)matching

Upon capturing hummingbirds, we photographed their bills against a 0.25-inch gridded notebook. We then used `ImageJ` to measure the length of the bill along the outer (top) surface of the bill using the 'line segment' feature after setting the scale using the gridded paper. To measure curvature, we utilized the methods of Temeles et al. (2009) in which we related the outer edge of the bill to a circle (Figure S2). The curvature of a circle is measured as the inverse of its radius. To estimate the radius of a circle with the same curvature as a bird's bill, we connected two points on the curved section of the bill, effectively drawing a chord on the circle (Figure S2). We then measured the angle between the chord and the line tangent (labeled $T$) to the bill at one end of the chord, known as the *angle of declension* ($\theta$ in Fig S2). Straight-forward trigonomentric identities then yield $\hat{r} = \frac{C/2}{\sin\theta}$, where $C$ is the length of the chord (mm) and $\theta$ is the angle of declension (radians). Thus, we use $\hat{r}^{-1}$ as an estimate of the curvature of a bill or flower.

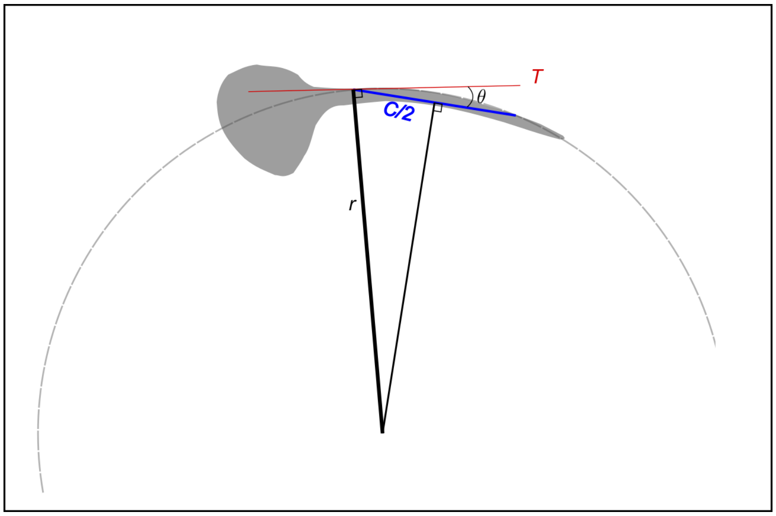


Figure S2: Relating the arc of the top of a bird's bill to a circle as in Temeles et al. (2009).

### Statistical methods

Let $y_{ijkl} \in \mathbb{N}$ be the number of pollen tubes scored in the $l^{\text{th}}$ flower from the $k^{\text{th}}$ plant of species $j = 1, .., s$, and let $i$ index the treatment $(i = 1, ..., g)$. We assume that $y_{ijkl} \overset{iid}{\sim} \text{Poisson}(\lambda_{ijk})$ for $l = 1, ..., n_k$. The model for the experiment can be written as

$$\log(\lambda_{ijk}) = \mu + \alpha_i + \beta_j + (\alpha\beta)_{ij} + \gamma_{k(j)}$$

where

- $\mu$ is the overall mean log-pollen tube rate,
- $\alpha_i$, $\sum_{i=1}^{g} \alpha_i = 0$, is the average deviation from the mean for flowers that received treatment $i$,
- $\beta_j$, $\sum_{j=1}^{s} \beta_j = 0$, is the average deviation from the mean for flowers of plant species $j$,
- $(\alpha\beta)_{ij}$, $\sum_{i=1}^{g} (\alpha\beta)_{ij} = \sum_{j=1}^{s} (\alpha\beta)_{ij} = 0$, is the species $\times$ treatment interaction effect,
- $\gamma_{1(1)}, \gamma_{2,(1)}, ..., \gamma_{K_1(1)}, \gamma_{1(2)}, ..., \gamma_{K_s(s)} \overset{iid}{\sim} \mathcal{N}(0, \sigma_\gamma^2)$, for the $k = 1, .., K_j$ plants of species $j$ and all species $j = 1, 2, ..., s$ are random effects for the plants that are nested within species.

For ease of defining the model in the `R` package `lme4` (Bates et al. 2015), we reparameterized the model to follow a regression parameterization (reference level for the aviary experiments was hand pollination for

*H. hirsuta* and hand pollination for *H. rostrata* for the nectar removal experiments). The model therefore reads

$$\log \boldsymbol{\lambda} = \mathbf{X}\boldsymbol{\beta} + \mathbf{Z}\boldsymbol{\gamma},$$

where $\mathbf{X}$ is a matrix of indicator variables indicating the species and treatment and $\mathbf{Z}$ is a matrix of indicator variables indicating the plant individual for the $i^{\text{th}}$ observation, $i = 1, ..., n$.
